# Visual acuity performance level is independent of locomotion

**DOI:** 10.1101/750844

**Authors:** Alex D. Swain, Eunsol Park, Zhang Yu Cheng, Nina Kowalewski, Angela Sun, Tessa Allen, Sandra J. Kuhlman

**Affiliations:** Program in Integrative Systems Biology, University of Pittsburgh; Department of Biological Sciences, Carnegie Mellon University; Carnegie Mellon Neuroscience Institute

**Keywords:** acuity, Go/No-go, locomotion, vision, perception

## Abstract

Locomotion has a global impact on circuit function throughout the cortex, including regulation of spatiotemporal dynamics in primary visual cortex (V1). The mechanisms driving state-changes in V1 result in a 2-3 fold gain of responsiveness to visual stimuli. To determine whether locomotion-mediated increases in response gain improve the perception of spatial acuity we developed a head-fixed task in which mice were free to run or sit still during acuity testing. Spatial acuity, ranging from 0.1 to 0.7 cycles/°, was assessed before and after 3-4 weeks of reward-based training in adult mice. Training on vertical orientations once a day improved the average performance across mice by 22.5 ± 0.05%. Improvement transferred to non-trained orientations presented at 45°, indicating that the improvement in acuity generalized. Furthermore we designed a second closed-loop task in which acuity threshold could be directly assessed in a single session. Using this design, we established that acuity threshold matched the upper limit of the trained spatial frequency; in two mice spatial acuity threshold reached as high as 1.5 cycles/°. During the 3-4 weeks of training we collected a sufficient number of stimulus trials in which mice performed above chance but below 100% accuracy. Using this subset of stimulus trials, we found that perceptual acuity was not enhanced on trials in which mice were running compared to trials in which mice were still. Our results demonstrate that perception of spatial acuity is not improved by locomotion.

## INTRODUCTION

Locomotion activates a powerful subcortical circuit that is associated with an increase in response gain in the visual cortex of mice (Ayaz et al., 2013; Bennett et al., 2013; Dipoppa et al., 2018; Erisken et al., 2014; Keller et al., 2012; Lee et al., 2014; Niell and Stryker, 2010; Pakan et al., 2016). During this state change, within the primary visual cortex (V1) the detection of weak stimuli is improved compared to rest (Mineault et al., 2016). The increased response gain has been shown to enhance neural encoding of visual stimuli (Dadarlat and Stryker, 2017; Mineault et al., 2016). These results raise the possibility that perception is enhanced during locomotion. Indeed, performance on perceptual visual tasks is improved during locomotion and the activation of the subcortical basal forebrain (Bennett et al., 2013; Goard and Dan, 2009; Pinto et al., 2013). Given that response gain is preferentially enhanced for higher spatial frequencies relative to lower spatial frequencies during locomotion (Mineault et al., 2016), together these results suggest that spatial acuity may be improved during locomotion relative to rest. On the other hand, during locomotion the retinal image can become blurred (Barlow and Olshausen, 2004) and it is possible that the enhanced representation of high spatial frequencies is necessary to counteract the loss of information that can occur during self-motion relative to a stationary environment. In the latter case locomotion would not be expected to improve spatial acuity. To distinguish between these two possibilities we designed a head-fixed task in which mice were free to run or sit still during acuity testing. We found that perceptual acuity was maintained but not improved during locomotion. Our results support the latter possibility. Performance on our head-fixed version of the acuity task was comparable to the more commonly used 2-alternative forced-choice visual water task (Hosang et al., 2018; Prusky and Douglas, 2004; Wang et al., 2016) in which mice are required to be in motion during acuity assessment. In addition, similar to the visual water task, we found that daily participation in the head-fixed acuity task resulted in improved spatial acuity.

## RESULTS

We developed a head-fixed task to assess perceptual acuity in mice. Mice were first trained to report the detection of vertically oriented static sine-wave gratings presented at a spatial frequency of 0.1 cycles/° (**Fig. 1**). We refer to this task as a ‘shaping’ task because it prepared the mice for the subsequent acuity assessment tasks. The task design followed a Go/No-go structure in which one of two stimuli (either a vertical grating ‘Go’ stimulus or an isoluminant gray screen ‘No-go’, stimulus) were presented with a fixed probability for the duration of the session. During ‘Go’ stimulus trials two possible behaviors were scored: one, the successful collection of the water reward, referred to as a ‘hit’, and two, a failure to collect the reward, referred to as a ‘miss’. Two possible behaviors were also scored on the ‘No-go’ stimulus trials: one, the successful withholding of a lick response for the entire duration of the trial, referred to as a ‘withhold’, and two, a failure to withhold a lick response referred to as a ‘false alarm’ (**Fig. 1A**). Our custom-built behavioral apparatus includes a photo-diode beam that the mice must first break with their tongue in order to retrieve the water reward on ‘Go’ trials. Because the mice are not cued by a drop of water appearing on ‘Go’ trials, latency to lick is an additional metric of performance that can be evaluated (**Fig. 1B**). Mice typically performed 150 trials in one session, and were not exposed to more than one session per day. After the first two days of training, false alarms were punished with a time out. Performance accuracy, an index ranging from 0 to 1, as well as d-prime continued to increase across sessions over the course of two-three weeks (**Fig. 1A,C**).

**Figure 1.**
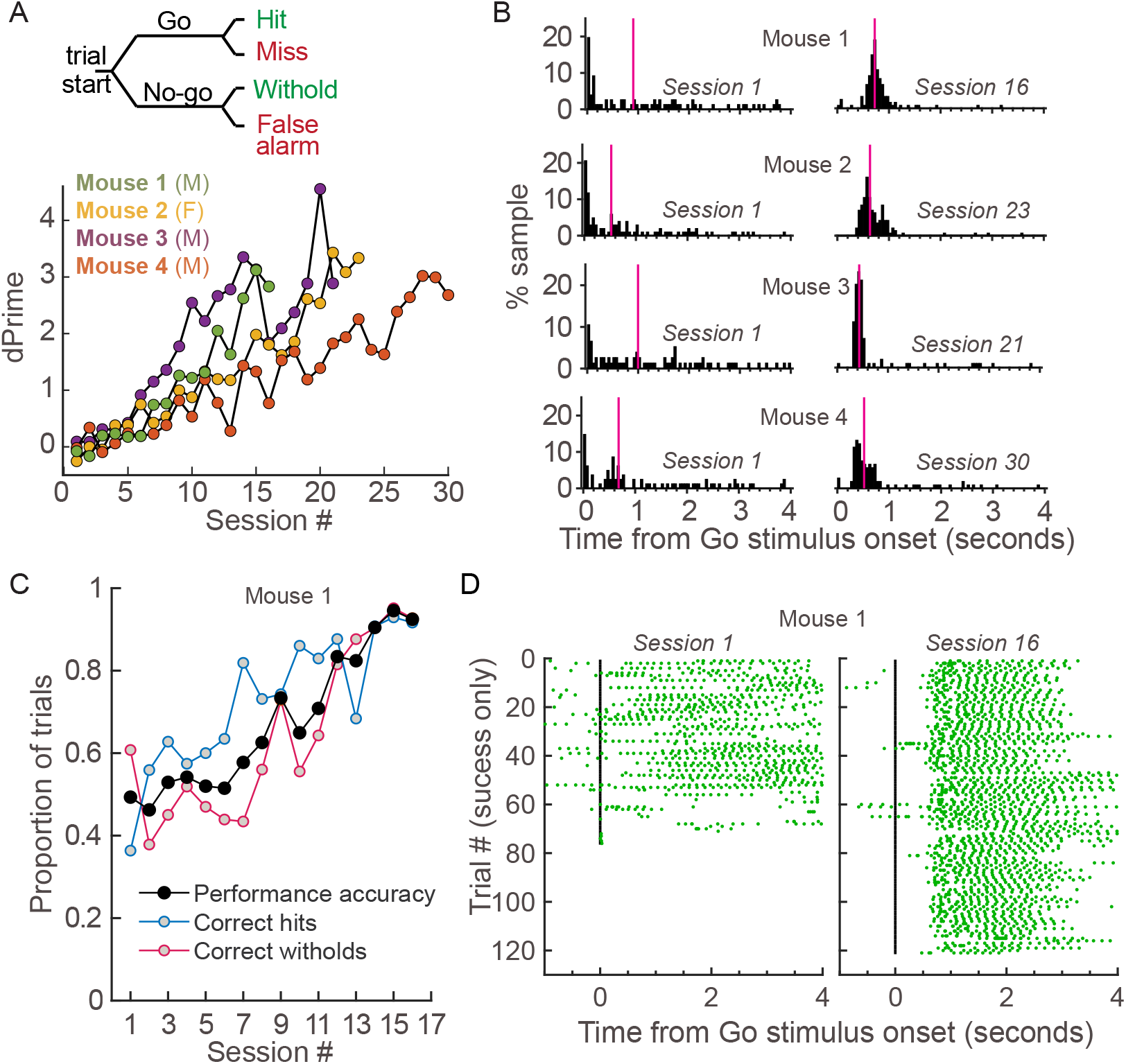
Head-fixed mice learn to report perception of vertically oriented gratings. A. Top, schematic of the Go/No-go discrimination task depicting four possible outcomes depending on stimulus type and behavioral response. Correct responses (green) and incorrect responses (red) are possible for both stimulus types. In this case, the ‘Go’ stimulus was a vertically oriented grating presented at 0.1 cycles/°, and the No-go stimulus was an isoluminant gray screen. Stimuli presentation was 4 seconds in duration followed by a two-second interval. False alarms were punished by a timeout. Mice performed 270-450 stimulus trials in one session per day. Bottom, discrimination accuracy of four mice, 3 males (M) and 1 female (F), reported as d-prime. B. Histograms of the first lick times across all ‘Go’ stimuli for the first and last session as indicated. Median lick time is indicated in pink. In all 4 mice the distribution of first lick times was significantly shifted to lower values on the last session compared to the first session (KS test p=1.48×10^−6^, p=1.86×10^−9^, p=1.8×10^−3^, p=6.26×10^−12^). *KS test, P<0.005 C. Performance of an example mouse increased across sessions, both hit rate and correct withhold rate (correct rejection) improve with training. D. Raster of lick times, aligned to the onset of the ‘Go’ stimulus (black vertical line), for an example mouse on the first and last sessions. Only ‘Go’ trials in which the mouse licked at least once are shown. Mice were free to lick more than once on ‘Go’ trials, however only received one water drop per ‘Go’ trial.

### Baseline spatial acuity performance in head-fixed mice is comparable to the visual water acuity task

In the shaping task, all mice achieved a d-prime of 3 or greater within 4 weeks. Latency to lick on ‘Go’ trials significantly decreased in all mice during this same time course (**Fig. 1B,D**). The median latency to lick decreased on average by 255 ± 104 ms (S.E.M.). Importantly, the withhold rate improved across sessions. When shaping was complete, the false alarm rate was lower than 13% for all mice. Based on this, we estimated that 23 ‘Go’ trials was the minimum number needed to yield sufficient statistical power to determine whether a specific spatial frequency was perceived in the acuity task.

To assess acuity, the above Go/No-go task was modified to include 6 different ‘Go’ stimuli, varying in spatial frequency (**Fig. 2A**). The design included 23 ‘Go’ stimuli for each spatial frequency tested, thus a single session was sufficient to evaluate acuity. In this design it is also possible to combine performance across sessions. For the results reported in this study two sessions were averaged. We found that the performance on the ‘Go’ stimulus presented at 0.2 cycles/°, a spatial frequency that the mice were not trained on during shaping, was indistinguishable from performance at 0.1 cycles/° (**Fig. 2B**). This demonstrates that the mice learned to lick in response to vertical gratings in general, and were capable of reporting the presence of gratings when perceived.

**Figure 2.**
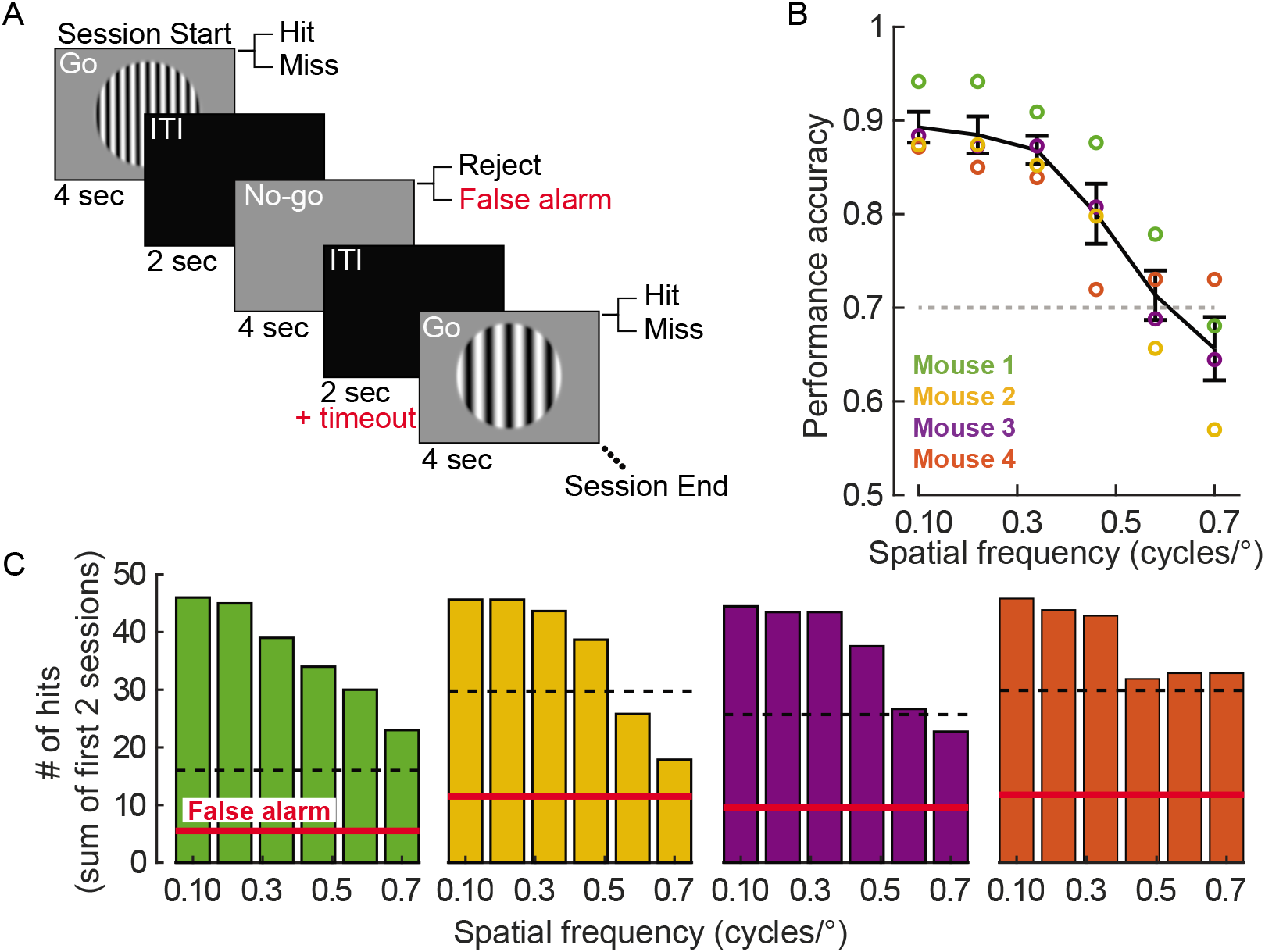
Performance decreases with increasing spatial frequency. A. Schematic of the acuity task design. One of 6 possible ‘Go’ stimuli were presented for a total of 23 times, randomly interleaved with ‘No-go’ stimuli. Each stimulus type has two possible behavioral outcomes and is followed by an inter-trial-interval (ITI). B. Performance accuracy decreased with increasing spatial frequency. Individual animals indicated by circle symbols. Each point is the average performance accuracy across the first two sessions. Dashed line indicates 0.7 performance accuracy. Error bars: SEM across animals. C. Hit rate, summed across the first two sessions and as such included forty-six ‘Go’ stimuli for each spatial frequency presented, decreased at higher spatial frequencies. False alarm rate, normalized to the total number of ‘Go’ stimulus presentations, is shown in red. The dashed line indicates the 95% confidence interval calculated using the false alarm rate which was averaged across the first two sessions.

In the acuity task, performance accuracy averaged across the first two baseline sessions was greater than 0.7 at 0.46 cycles/° for all mice, and greater than 0.7 for half of the mice at 0.58 cycles/°, and less than 0.7 for all mice except one at 0.7 cycles/°. These performance values are consistent with values obtained using the visual water task (Hosang et al., 2018; Prusky and Douglas, 2004; Stephany et al., 2018; Wang et al., 2016). In addition to performance accuracy, hit rate relative to the false alarm rate can be a useful indicator of performance when the false alarm rate is low. To assess whether the hit rate was above chance, a 95% confidence interval was generated based on the false alarm rate. Spatial frequencies for which the hit rate was above the 95% confidence interval were considered to be significant (**Fig. 2C**). We found that this metric was generally consistent with the 70% performance cut-off used to interpret performance accuracy in **Figure 2B**, however, there was a trend for performance accuracy values >0.68 to be scored as significant when using the hit rate method. Overall, the hit rate analysis validates using the 0.7 cut-off for performance accuracy, but does indicate that the 0.7 cut-off is conservative.

### Perceptual acuity is not enhanced by locomotion

Next we determined whether perceptual acuity for static gratings was improved during locomotion (**Fig. 3**). To do this, we exposed mice to at least 20 sessions of the acuity task to collect enough data for analysis. Individual mice exhibited variability in their preference for running, ranging from 13 to 70% of the trials (**Fig. 3A**). For this analysis we considered only trials in which mice had a hit rate of 16-22 hits/ session to eliminate the confound of floor or ceiling effects. We found that the withhold rate was negatively correlated with locomotion across mice (**Fig. 3B**). However, taking into account individual differences in mice, this effect appears to be driven by variation across mice rather than an effect present within the individual. We found no correlation between the withhold rate and the amount of locomotion (Spearman rank correlation p values, mouse #1 through 4: 0.83, 0.73, 0.43, 0.78). Similarly, we found that hit rate was not enhanced during locomotion (**Fig. 3C**). Neither a correlation across animals nor within animals was detected. To provide additional support for these results, we included all sessions, regardless of performance, and examined whether locomotion had an impact on performance accuracy (**Fig. 3D**). We found that there was only one case in which locomotion had a significant effect on performance; locomotion significantly decreased performance accuracy in mouse #2 for the 0.46 cycles/° ‘Go’ stimulus. In summary, our analyses were robust enough to determine whether locomotion improved acuity, and we found that perceptual acuity assayed using static gratings was not enhanced during running.

**Figure 3.**
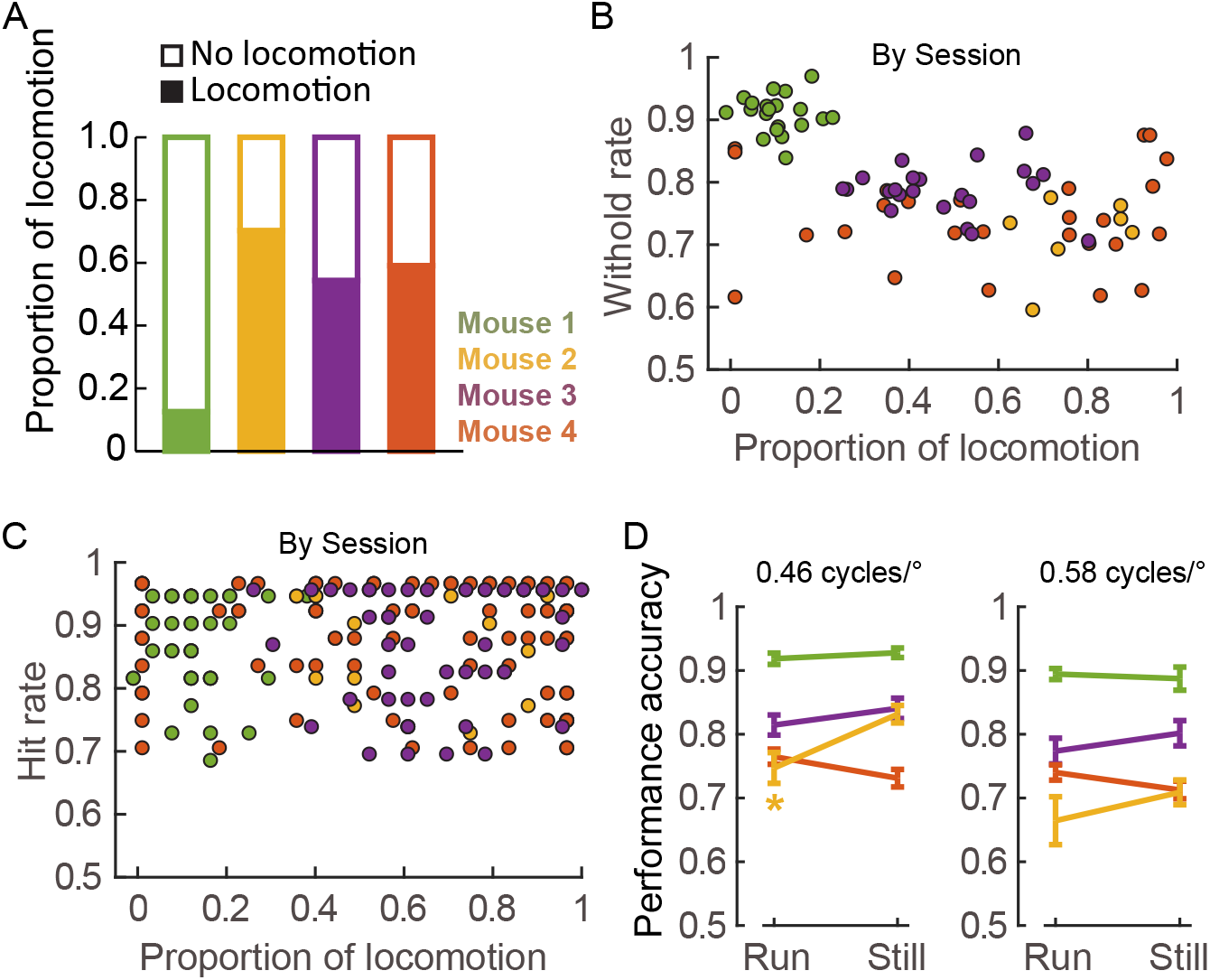
Perceptual acuity is not enhanced by locomotion. A. Proportion of trials in which animals exhibited locomotion during the course of acuity training. B. Withhold rate of individual mice (indicated by color) relative to the proportion of locomotion trials. Each data point represents a single session. All sessions for which locomotion data was available are plotted. Any spatial frequency meeting this criterion is included. Across the population, there was a negative correlation between withhold rate and locomotion (Spearman correlation r= −0.58, p= 3.8×10^−8^). C. Hit rate of individual mice (indicated by color) relative to the proportion of locomotion trials. Individual data points are offset by 0.025% to visualize overlapping points. Spearman correlation considering all data points was not significant (p=0.46). Only trials in which hit rate was greater than 16 and less than 20 were included, all spatial frequencies were considered. D. Performance across all trials for ‘Go’ stimuli presented at 0.46 and 0.58 cycles/°, individual mice plotted as indicated. In one mouse performance was significantly higher in the still condition (* t-test, p=0.0183).

### Perceptual acuity can improve up to 1.5 cycles/° with training

During the course of collecting the above data, we noted that after 3 days of training there were indications that perceptual acuity was improving. Therefore, we analyzed the extent to which extended training improved acuity (**Fig. 4**). Regression analysis revealed that all mice significantly increased performance accuracy on the ‘Go’ stimulus presented at 0.7 cycles/° (**Fig. 4A**). Furthermore, for spatial frequencies that require the cortex for perception [frequencies greater than 0.3 cycles/° (Prusky and Douglas, 2004)], we found that performance accuracy improved (**Figs. 4B,C**), and that there was a significant decrease in the latency to lick (**Fig. 5**).

**Figure 4.**
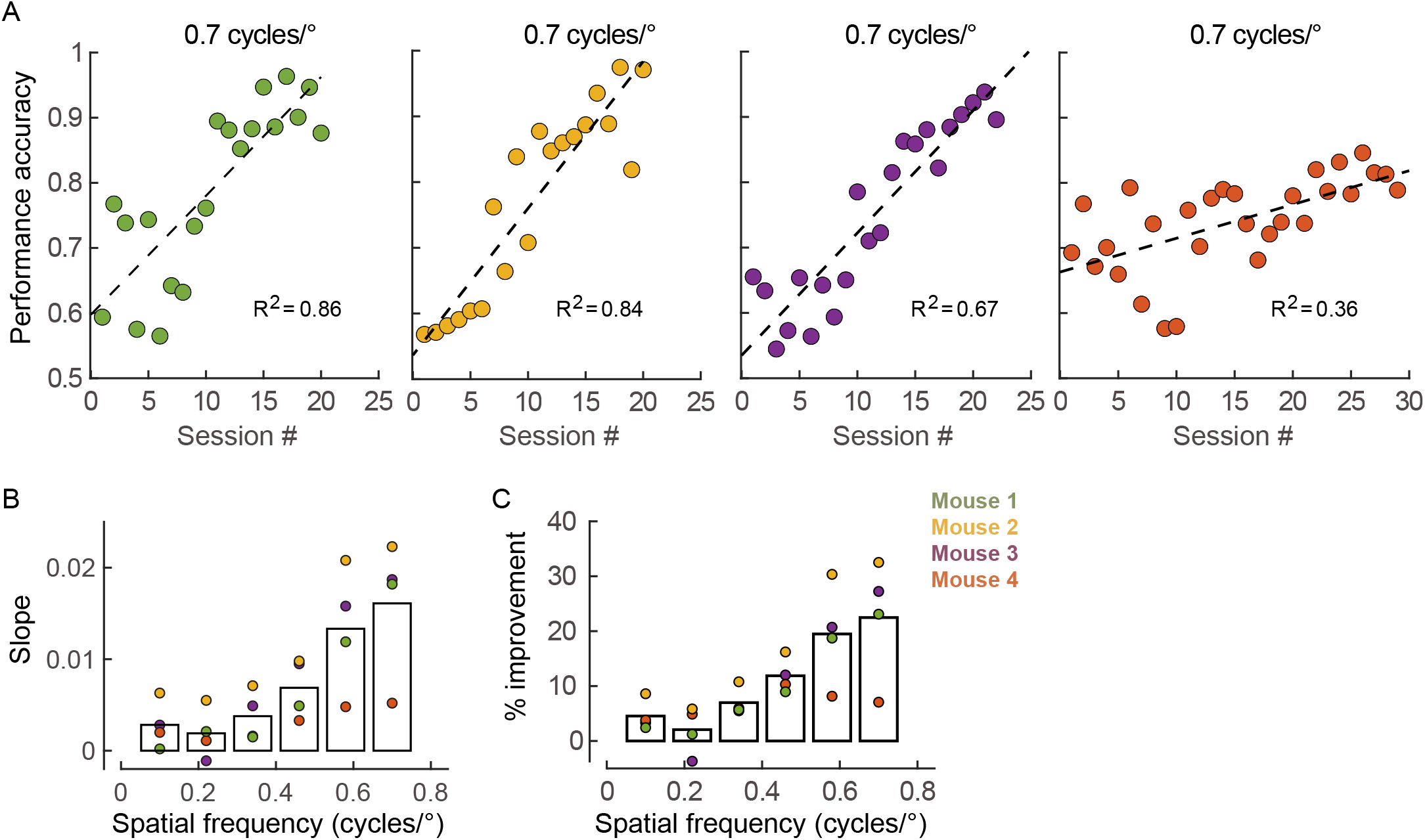
Perceptual acuity is improved with training. A. All mice increased performance accuracy across sessions (linear regression, p= 9.24×10^−16^, p= 1.93×10-23, p=2.74×10^−12^, p=1.0×10^−13^, mouse#1-4 respectively). B. Slope of the regression fit across training sessions. Data points represent individual mice as indicated, and the mean across mice is indicated by bars, calculated separately for each spatial frequency. C. Percent improvement, the average of performance accuracy for the first two sessions compared to the last two sessions.

To determine whether improvement generalized to non-trained orientations, the perceptual acuity of mice was probed for gratings presented at an orientation of 45° (**Fig. 5**). We found that for all spatial frequencies greater than 0.3 cycles/°, performance accuracy was significantly higher after training at an orientation of 0° and 45° compared to baseline assessment tested at an orientation of 0° (**Fig 5A**). Similarly, the median latency to first lick across ‘Go’ trials was lower for both orientations after training (**Fig. 5B**). In all mice, the distribution of latencies to lick was significantly shifted towards lower values after training for both orientations (**Fig. 5C**).

**Figure 5.**
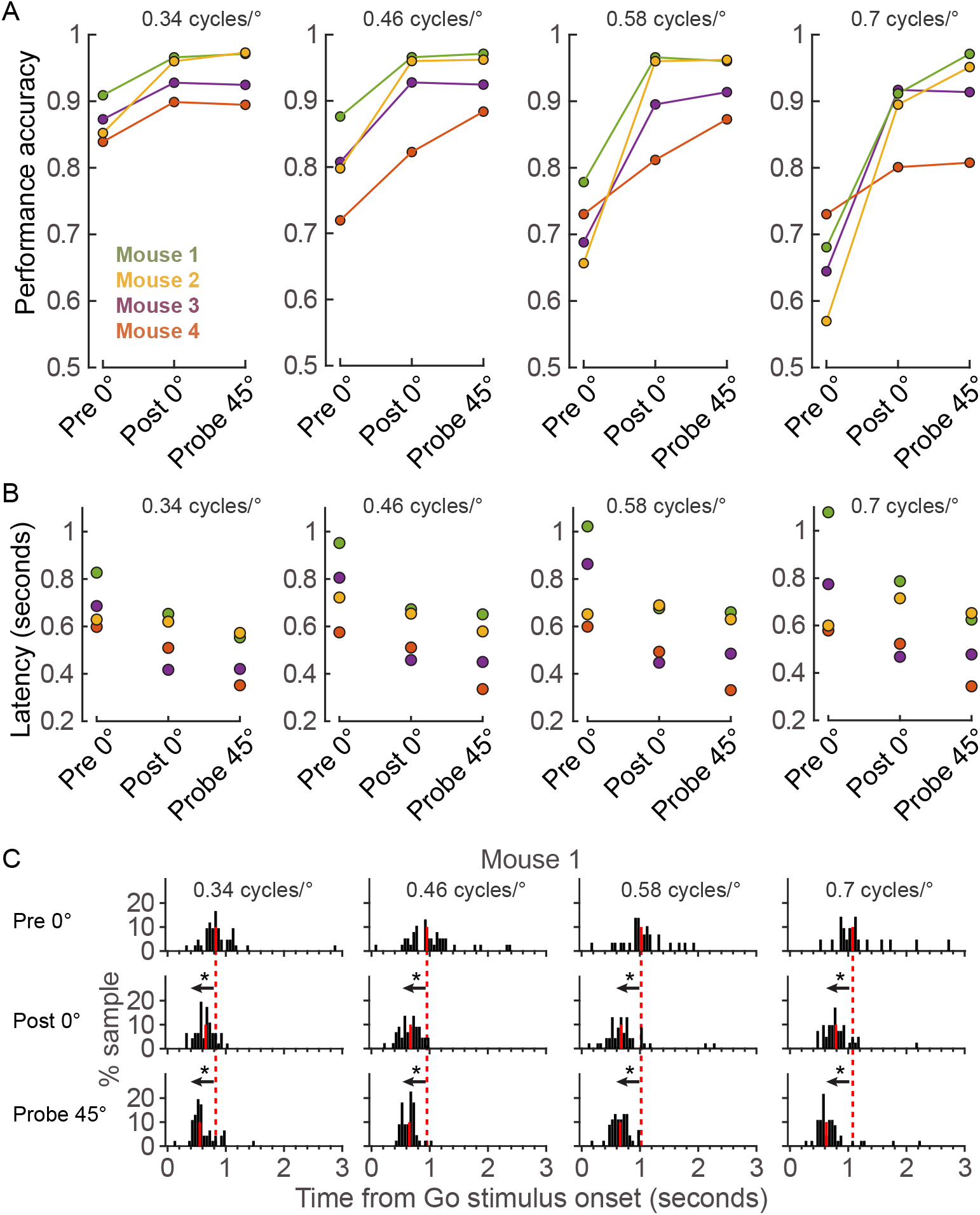
Acuity improvement generalizes to a non-trained orientation. A. Average performance accuracy across two sessions prior to training (Pre), after training (Post) and post-training probe (Probe) for each mouse. Angle and spatial frequency (by column) of ‘Go’ stimulus presentation as indicated. B. Latency to first lick across two sessions prior to training (Pre), after training (Post) and posttraining probe (Probe) for each mouse. Angle and spatial frequency (by column) of ‘Go’ stimulus presentation as indicated. C. Histograms from one example mouse of the first lick times across all ‘Go’ stimuli prior to training (Pre, row 1), after training (Post, row 2) and post-training probe (Probe, row 3). Spatial frequency (by column) of ‘Go’ stimulus presentation as indicated. The distribution of first lick times was significantly decreased for both the Post and Probe conditions compared to the Pre condition (KS test p values, Pre vs Post increasing spatial frequencies: 9.4×10^−6^, 3.9×10^−6^, 4.89×10^−5^,7.7×10^−3^; Pre vs Probe increasing spatial frequencies: 8.8×10^−5^, 7.5×10^−5^, 4.5×10^−4^, 4.2×10^−3^). *p<0.01

The acuity assessment paradigm described above is useful to determine spatial acuity for a specific spatial frequency. However, in one session it is not possible to determine the acuity threshold of an individual animal using this paradigm. We developed a second acuity assessment paradigm in which the threshold can be assessed in a single session for a given animal (**Fig. 6**). A block design was used in which two different spatial frequencies (SF_1_ and SF_2_) were presented as ‘G’o stimuli in addition to a gray screen ‘No-go’ stimulus. In this paradigm hit rate on a given block is used to determine the next spatial frequencies to be presented; as such this is a closed-loop design in which the behavioral response of the animal dictates the next stimulus to be tested. Threshold is determined by identifying the spatial frequency that SF_1_ and SF_2_ converge to. To demonstrate that the closed-loop block design could be used to determine acuity threshold, two of the mice described above (mouse #2 and #3) were trained for additional sessions, using the extended training acuity paradigm. During extended acuity training, these mice were exposed to spatial frequencies as high as 1.5 cycles/°. After extended training, mice were subjected to the acuity threshold assay (**Fig. 6A**). We found that perceptual acuity determined using the threshold assay closely matched the highest spatial frequency that the mice were exposed to during extended training. Eight blocks presented in one session were sufficient for SF_1_ and SF_2_ to converge (**Fig. 6B**).

**Figure 6.**
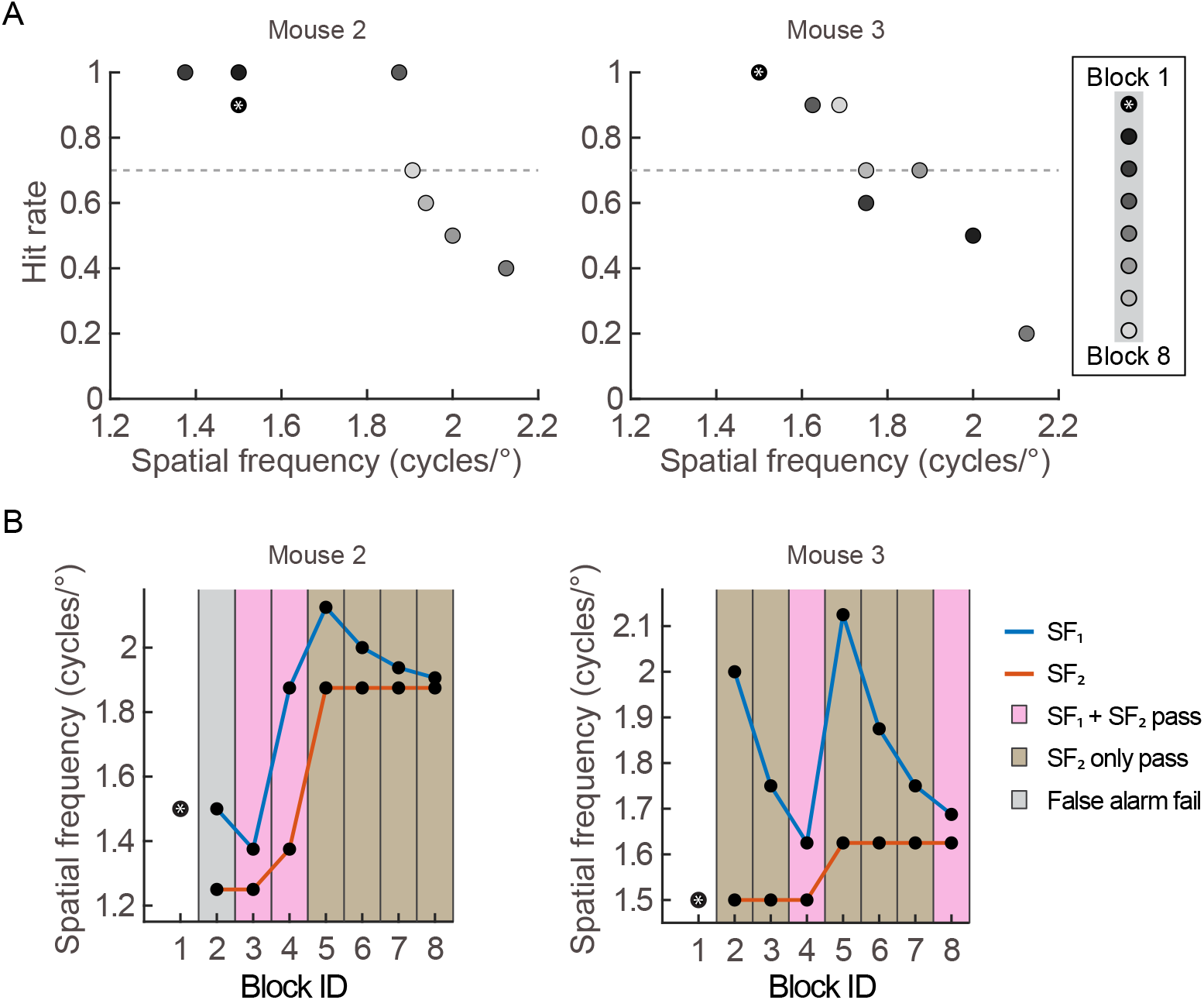
Acuity threshold limit matches the training stimuli. A. Spatial frequency of ‘Go’ stimulus versus hit rate for two mice as indicated. Dashed line indicates 0.7 hit rate. B. Progression of spatial frequencies presented over the course of the closed-loop task for two mice as indicated.

## DISCUSSION

The mouse is an established model for developing new methods to enhance vision in amblyopic animals (Gordon and Stryker, 1996; Hensch and Quinlan, 2018; Stryker and Lowel, 2018). In this line of research there are two goals: one, to restore ocular dominance plasticity and two, to improve acuity in the eye-specific pathway that was deprived of normal sensory experience during development. In recent years much progress has been made in identifying the molecular components and signaling pathways that can be activated to restore ocular dominance plasticity. However, there is a gap in our understanding of how to improve spatial acuity after responsiveness is restored. Indeed, it was recently demonstrated that ocular dominance plasticity and the development of acuity are mechanistically distinct and represent dissociable processes (Stephany et al., 2018). The gap in knowledge exists in part because ocular dominance is straightforward to classify for the majority of neurons recorded from during an experiment, whereas determining the spatial cut-off frequency for a neuron requires a more extensive range of stimuli to be tested and is therefore more difficult to determine. The ability to assay the response properties of neurons at a high density using improved calcium imaging methods circumvents this issue (Jeon et al., 2018). To take full advantage of this approach, there is a need to develop perceptual assays of spatial acuity that are compatible with high density recording. To that end, here we describe two different paradigms for assessing perceptual acuity in head-fixed mice. Importantly, we demonstrated that mice can learn the association between grating stimulus and reward at low spatial frequencies and immediately transfer the association to higher spatial frequencies.

To compare our task with the established 2-alternative forced choice visual water task in which mice are placed in a water chamber containing two arms (Prusky and Douglas, 2004), we first assessed spatial acuity in the head-fixed task. In the visual water task mice must select the arm in which a grating stimulus is presented (versus a gray screen) and swim to a hidden escape platform present in the same arm as the grating stimulus. This task has the advantage that mice do not need to be trained on the apparatus, their innate behavior is to escape from water. Similar to our head-fixed task, mice do need to learn the association of the grating stimulus with reward. We found that baseline acuity measurements were comparable between the two tasks. These results establish that our Go/No-go acuity task is a valid method to assess visual acuity in mice. Furthermore, we found that similar to the visual water task (Hosang et al., 2018; Wang et al., 2016), reinforced training in adult mice improved acuity by >30%. The improvement generalized to orientations that the animal was not trained on, confirming that actual acuity was improved. In standard rearing conditions, perceptual acuity increases during postnatal development and plateaus at approximately 0.45-0.55 cycles/° by age postnatal day (P) 45 (Prusky and Douglas, 2004; Stephany et al., 2018). The finding that spatial acuity can be improved past the critical period for ocular dominance is consistent with the observation that ocular dominance plasticity and the development of acuity are two dissociable processes.

Five parameters can be measured in our head-fixed acuity task: d-prime, latency to lick, performance accuracy, hit rate, and withhold rate. We recommend using performance accuracy as the primary metric of performance in the acuity testing paradigm. Performance accuracy weighs correct withholds equally with correct hits. Given mice have a tendency to impulsively lick in Go/No-go tasks, it is important to use a metric that fully accounts for No-go trials in which the animal makes an incorrect response. In the case of the acuity threshold task, given that there are only 40 ‘No-go’ stimuli per block and our mice all had low false alarm rates, we found that hit rate was the most useful metric of performance.

Accumulating evidence raises the possibility that perceptual acuity is enhanced during locomotion. To directly determine whether this is the case, we used our head-fixed paradigm in which it is possible for mice to retrieve their reward either in a state of locomotion or in a quite state. We found that locomotion did not enhance perceptual acuity. Importantly, hit rate was not impacted by locomotion. Consistent with this, performance accuracy was not altered by locomotion, except one mouse performed slightly better when still at a spatial frequency of 0.46 cycles/°. We did find a negative correlation between the withhold rate and locomotion across animals. This may be an indication that when running the animal has a harder time refraining from impulsively licking on a ‘No-go’ stimulus trial. However, we did not see a correlation between withhold rate and locomotion within an individual animal across sessions. Taken together, our interpretation is that acuity perception is not lower on trials in which there is locomotion compared to still trials for a given animal. However our results do raise the possibility that the absolute performance accuracy of an individual mouse may depend on locomotion preference in the head-fixed acuity assessment task. Because the mouse is a useful model to test the impact of manipulating molecular pathways on perception, it will be important to take this into account when interpreting the impact of targeted genetic mutations or molecular perturbation on visual acuity measurements when using this task. Our results are consistent with previous studies demonstrating that although frequently associated with one another, locomotion and arousal are two distinct states and differentially impact the encoding of visual stimuli (Reimer et al., 2014; Vinck et al., 2015).

We used static gratings in the present study. The extent to which our results apply to stimuli containing motion, including drifting gratings, remains to be determined. The mouse visual system contains two parallel processing pathways that resemble dorsal and ventral streams found in primates (Smith et al., 2017). The ventral stream is necessary for the detection of form, while the dorsal stream contributes to the perception of motion in visual stimuli. It has been observed that locomotion enhances at least two aspects of the representation of information at the neural level within the primary visual cortex (V1) when mice view drifting gratings (Mineault et al., 2016). Furthermore, detection of low contrast drifting gratings is enhanced during locomotion(Bennett et al., 2013). Thus, it is possible that our results apply specifically to high-contrast static gratings.

In summary, our results are consistent with the possibility that increased response gain to static images serves to offset motion-induced sources of noise such as retinal blur rather than enhancing perceptual acuity. The locomotion-induced improvement in perception observed in previous studies using low-contrast motion stimuli (Bennett et al., 2013; Pinto et al., 2013) likely involves the enhancement of processes other than the detection of spatial frequency per se.

## METHODS

Experimental procedures were compliant with the guidelines established by the Institutional Animal Care and Use Committee of Carnegie Mellon University and the National Institutes of Health. Experimental mice (1 female and 4 males) were generated from the following cross: ErbB4 floxed heterozygote (MMRC, stock number 010439-UCD), PV-cre homozygote (Jackson Laboratories, stock number 005628) X ErbB4 floxed heterozygote. 50% of the offspring carried one functional copy of the ErbB4 allele and 25% of the offspring carried no functional copies of the ErbB4 allele; neither of the mutant genotypes was used in this study. Mice were group housed on a reverse light cycle in standard cage conditions that included nesting material and a domed hut. Food was provided *ad libitum*.

### Head-post surgery and water restriction

As described (Feese et al., 2018; Pafundo et al., 2016), mice (age ≥ p28) were anesthetized with isoflurane (3% for induction, 1-2% for maintenance; Patterson Veterinary; 07-893-1389). Skin covering the dorsal surface of the skull was removed, bone scuffed and a layer of cyanoacrylate was applied. A custom-made stainless-steel bar was glued to the right side of their skull and secured with dental cement (Lang Dental; 1404R and 1220pnk). The skull surface was sealed with additional dental acrylic. One dose of Carprofen (Rimadyl; Patterson Veterinary, cat# 07-844-7425) was injected subcutaneously (0.25 ml volume, 0.5 mg/ml) prior to surgery, and a second dose 24-hours after the surgery. 48-hours after surgery mice began water-restriction, receiving 750μl per day (their daily ration) for 8 to 10 days until body weight stabilized. Once stabilized, behavioral tasks were initiated. The weight and health of all water-restricted mice were monitored daily.

### Behavioral apparatus and visual stimuli

A 3D-printed custom acrylic lickport was used to record licks and to deliver water rewards. It was placed in front of the mouse within reach of the mouse’s tongue. Licks were recorded by the tongue breaking an infrared light path between LED optical switch photodiodes (Vishay Semiconductors; TCZT8020-PAER). Water rewards were delivered by gravity using a 3-port solenoid valve that opened for a defined period of time following a beam-break on ‘Go’ trials (Lee Company; LHDA1231115H). Water rewards were delivered into the lickport through a 0. 02-inch-diameter stainless steel tube. In order to prevent the pooling of water on the lickport, excess or pooled water was pumped away from the steel reward tube by plastic tubing using a peristaltic pump (Fisherbrand; 70730-064). An mBed microcontroller processor (mBed 1768 Demo Board; Mouser 7711-OM11043598) and custom scripts were used to schedule ‘Go’ and ‘No-go’ stimuli, sample lick times at a frequency of 10 Hz, and gate release of water rewards.

The volume of the water delivered per beam beak was controlled by the duration that the solenoid valve was open. For the acclimation task, water drop size was ~3-4μL, for Go/No-go tasks the rewarded volume was ~6-7μL. During behavior, mice were mounted on stainless steel bar positioned over a Styrofoam ball floated on a cushion of air. The screen (Dell; 30”, 2560×1600 resolution; 9TDTX) was positioned 25cm away from the mouse in front of the right eye, angled at 50° with respect to the midline of the animal.

Rewarded ‘Go’ sinusoidal grating stimuli were presented at a diameter of 60° against a gray background at 100% contrast; edges were smoothed with a Gaussian blur (σ=10) to eliminate sharp edges. The ‘No-go’ stimulus was an isoluminant gray screen. Stimuli were generated using Psychophysics Toolbox (http://psychtoolbox.org) in Matlab (Mathworks, Boston, MA).

Mice were mounted into the apparatus by briefly anesthetizing (≤ 2 minutes, 3% induction) with isoflurane and attached to a steel bar. The steel bar was then inserted into a holder to position the animal over the center of the Styrofoam ball. Mice were given 5 minutes to balance prior to initiating behavioral tasks.

### Lickport acclimation

The goal of lickport acclimation is to familiarize mice with obtaining water from the lickport. A ‘Go’ stimulus (vertically orientated grating at a spatial frequency of 0.1 cycles/°) was presented to the mouse for a variable amount of time. The stimulus duration was manually controlled by the user, according to the following rule: the ‘Go’ stimulus appeared until the mouse broke the beam and received ~2-5 drops of water, and at that time was switched to the No-go stimulus until the mouse stopped licking for ~60 seconds. This sequence continued until the mouse received the daily ration or stopped licking for 10 minutes. Typically on the second day of acclimation mice received the daily ration within 30 minutes. If on the first session less than 500 μl was received, the mouse was supplemented to 500 μl. On all subsequent sessions the mouse was supplemented to 700 μl, a volume slightly below the daily ration. Mice performed one session per day.

### Go/No-go tasks

Stimulus trials were separated by an inter-trial-interval of 2 seconds during which a black screen was presented. For every ‘Go’ trial that the mouse licked, one water drop was delivered. False alarms (licking on a ‘No-go’ stimulus) were punished with a timeout: the black screen inter-trial-interval duration was extended for 1 to 8 seconds, depending on the mouse (some responded well to 1 second, while others required a longer duration). If the mouse licked at any point during the timeout, the timeout duration was reset and triggered again. Timeout was not used on the first two days of shaping. Misses (a failure to lick on the ‘Go’ stimulus) were not punished. Mice performed one session per day. Four variants of the Go/No-go task were used:

#### 1. Shaping

The goal of the shaping task was to teach mice to lick on a rewarded stimulus, and withhold licks on a non-rewarded stimulus. If the mice did not receive their daily ration, they were not supplemented for that session. In the case mice did not receive the daily ration for two days in a row they were supplemented to 700μL on the second day. Stimuli were 4 seconds in duration; the Go stimulus was a vertical oriented grating of 0.1 cycles/°. The stimulus schedule was semirandom. Three Go stimuli could not appear in a row. The proportion of ‘Go’ trials to ‘No-go’ trials was modified across sessions. The probability of a ‘Go’ stimulus was set to 0.46 on the first day of shaping, and decreased to 0.25 on the final shaping session. Mice were shaped until they achieved 90% or greater performance for 2 to 3 sessions. False alarm rate was defined as the number of ‘No-go’ trials within a session in which there was at least one lick. One mouse (mouse#3) achieved this criteria, but do due to a break in training immediately following reaching criteria was trained for an additional 6 days until it reached 3 consecutive sessions of >90% performance. Performance for the shaping task was calculated as follows:

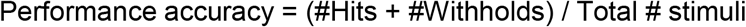

Where a hit is at least one lick on a Go stimulus, and a withhold is when the mouse did not lick on a No-go stimulus. Hit rate and withhold rate were calculated as follows:

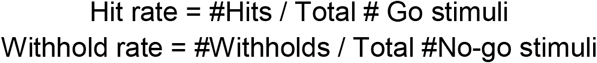

#### 2. Acuity Testing

There were six ‘Go’ stimuli presented at six different spatial frequencies at fixed angle for a given session. ‘Go’ stimuli were presented with a probability of 0.30. Each ‘Go’ stimulus was presented 23 times, as such there were a total of 138 possible water-rewarded trials in one session. Spatial acuity was determined by averaging their performance at each of the six spatial frequencies for the first two sessions of testing. Including 46 trials rather than 23 increased the statistical power when comparing acuity between animals. Performance was calculated for a given spatial frequency ‘Go’ stimulus (S) as:

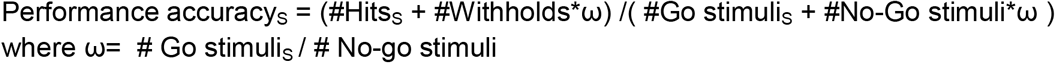

where ω= # Go stimuli_S_ / # No-go stimuli ω is a weight factor that is used in order to account for the difference in the number of No-go stimuli compared to a given ‘Go’ stimulus. If a mouse received less than 600μL on a given session, it was supplemented to 600 μl.

#### 3. Extended acuity training

Mice were exposed to extended acuity testing for 18 to 27 sessions after the initial 2 days of baseline assessment. Initially, the range of spatial frequencies used to assess spatial acuity was 0.1, 0.22, 0.34, 0.46, 0.58, 0.70 cycles/°. In the case of three of the four tested mice, their performance at high spatial frequencies [0.58, 0.7 cycles/°] improved, as such we replaced the 0.22 cycles/° Go stimulus with 0.82 cycles/°. The final spatial acuity presented in Figures 4 and 5 was assessed as the performance for each spatial frequency averaged across the last two days of extended training. Two of the four mice continued to receive extended training for 10 or more sessions and were exposed to spatial frequencies as high as 1.5 cycles/° by replacing the lowest spatial frequency greater than 0.1 cycles/° that the mouse was exposed to with the next highest spatial frequency the mouse had not yet been exposed to, in steps of 0.12 cycles/°. If a mouse received less than 600μL on a given session, it was supplemented to 600μL. Performance accuracy was calculated as above.

#### 4. Acuity Threshold Determination

The goal of the threshold task was to determine the limit of spatial acuity for an individual mouse in one session. A block design was used in which two different spatial frequencies (SF_1_ and SF_2_) were presented as ‘Go’ stimuli in addition to a gray screen No-go stimulus, to identify the spatial frequency at which performance on the two ‘Go’ stimuli converged above a user-defined threshold. In this study we chose the threshold to be a hit rate of ≥ 0.8 and false alarm rate of ≤ 0.3. In the case performance on both SF_1_ and SF_2_were above threshold, in the next block SF_1_ (the higher of the two) was increased by 0.5 cycles/° and SF_2_ was increased to the former level of SF_1_. In the case performance of SF_2_ only was above threshold, SF_1_ was decreased by half of the difference between the former SF_1_ and SF_2_. In the case performance of both SF_1_ and SF_2_ were below threshold, SF_1_ was decreased by half of the difference between the former SF_1_ and SF_2_ (this specific case did not occur in the experiments presented here). In addition, in the case performance on both SF_1_ and SF_2_ were above threshold for two consecutive blocks, SF_1_ was increased by half of the difference between SF_1_ and SF_2_, and SF_2_ was increased to the former level of SF_1_. All trials were randomized.

The task was initiated (Block 1) by presenting a single ‘Go’ stimulus for 10 trials, and a No-go stimulus for 20 trials. If the false alarm rate was ≤ 0.3 and the hit rate was ≥ 0.8, SF_2_ was set to the spatial frequency used for initiation and SF_1_ was set to 0.5 cycles/° higher than SF_2_. If these criteria were not reached, SF_1_ was set to the spatial frequency used for initiation and SF_2_ was set 0.25 cycles/° lower than the spatial frequency used for initiation.

### Locomotion analysis

Three different behavioral rigs were used to test the acuity and obtain motion analysis information for the mice, and each mouse stayed on the same behavioral rig for the duration of its testing. Each Styrofoam ball painted with small black dots around the ball to facilitate tracking. Locomotion information was obtained using a Keyence LV-NH32 sensor head spot and LV-N11MN amplifier; this system outputs reflectance values as analog voltages (0 – 5V). Voltage signals were digitized using an Arduino (10-bit); sampling rate was 1KHz (1ms). The difference in reflectance between each sample was calculated and binned for 100ms. Due to minor differences between the rigs and the balls, a motion threshold was defined for each of the three behavioral rigs. The threshold is a value that defines if mouse was, or was not, in a state of motion. For two of the rigs, the threshold value for motion was defined manually: The threshold was obtained by manually curating the responses of a mouse during an acuity session. Users defined each trial as either “No-Run No-Go”, “Run No-Go”, “No-Run Go”, and “Run Go”. For one of the four mice, the manual curation was unable to be performed. Therefore, a distribution of the motion data across all trials in each session was created, and found to be bimodal. The threshold was defined as point of least overlap between the two distributions. We then estimated if the mouse was, or was not, in a state of motion just prior to generating a response. On trials when the mouse licked, the one-second preceding the lick time point was used as a window for estimating motion. In that one-second window, 10 motion values were present. If 80% or more of the motion values were above the motion threshold, the mouse was considered to be in a state of motion for that specific trial. On trials when the mouse did not lick, an estimation of their motion status was obtained. This was done by obtaining the average lick time and standard deviation of the mean for each stimulus type. A time window was established for each trial that the mouse did not lick, defined as:

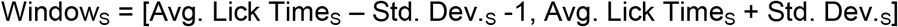

If 80% or more of the motion values were above the established motion threshold in that time window, the mouse was defined as being in a state of motion for that particular trial.

In order to establish the effect of motion on hit rates across sessions during extended acuity training we only considered hit cases where the mouse was performing above chance (that is, had at least 16 hits), but also did not perform perfectly (that is, had less than 23 hits). From these selected stimulus cases in a given session, the hit rate proportion for the ‘Go’ stimulus cases and motion was determined. A similar method was used to establish the relationship between withhold rate and motion. In this case, however, every session had a single withhold rate and a single motion proportion because there was only one type of ‘No-go’ stimulus with no selection criteria.

### Statistics

In cases the data were not normally distributed, non-parametric tests were used. Error bars report ± SEM.

## ACKNOWLEDGEM ENTS

We thank Brian Jeon for developing the behavioral apparatus and advice on collecting behavioral data, and Patricia Stan for help in designing the acuity threshold task.

## AUTHOR CONTRIBUTIONS

S.J.K., A.D.S., and E.P. designed the experiments. A.D.S., E.P., Z.Y.C., N.K., A.S., and T.A. collected the data. S.J.K. and A.D.S. wrote the manuscript.

